# Investigating an in-silico approach for prioritizing antidepressant drug prescription based on drug-induced expression profiles and predicted gene expression

**DOI:** 10.1101/2020.04.22.054940

**Authors:** Muhammad Shoaib, Edorado Giacopuzzi, Oliver Pain, Chiara Fabbri, Chiara Magri, Alessandra Minelli, Cathryn M. Lewis, Massimo Gennarelli

## Abstract

In clinical practice, antidepressant prescription is a trial and error approach, which is time consuming and discomforting for patients. This study investigated an in-silico approach for ranking antidepressants based on their hypothetical likelihood of efficacy.

We determined the transcriptomic profile of citalopram remitters by performing a transcriptomic-wide association study on STAR*D data (*N =1163*). The transcriptional profile of remitters was compared with 21 antidepressant-induced gene expression profiles in five human cell lines available in the connectivity map database. Spearman correlation, Pearson correlation, and the Kolmogorov Smirnov test were used to determine the similarity between antidepressant-induced profiles and remitter profiles, subsequently calculating the average rank of antidepressants across the three methods and a p-value for each rank by using a permutation procedure. The drugs with the top ranks were those having high positive correlation with the expression profiles of remitters and they may have higher chances of efficacy in the tested patients.

In MCF7 (breast cancer cell line), escitalopram had the highest average rank, with an average rank higher than expected by chance (*p=0.0014*). In A375 (human melanoma) and PC3 (prostate cancer) cell lines, escitalopram and citalopram emerged as the second highest ranked antidepressants, respectively (*p*=0.0310 and 0.0276, respectively). In HA1E (kidney) and HT29 (colon cancer) cell types, citalopram and escitalopram did not fall among top antidepressants.

The correlation between citalopram remitters’ and (es)citalopram-induced expression profiles in three cell lines suggests that our approach may be useful and with future improvements it can be applicable at the individual level to tailor treatment prescription.

## Introduction

Major Depressive Disorder (MDD) is a primary health issue and the third leading cause of disability in adolescents and young adults, while the second leading cause of disability in middle aged adults on a global scale (1). According to the World Health Organization, more than 264 million people are living with depression worldwide. This heavy disease burden is partly due to the complex pathogenic mechanisms of MDD, the inter-individual heterogeneity of antidepressant response and the lack of reliable response predictors (2).

Antidepressant (AD) choice in MDD is based on prescription guidelines and prior clinical experience, but the lack of reproducible predictors of AD response makes it a ‘trial and error’ approach which can take up to several weeks or months and a number of treatment changes before symptom remission is achieved. The availability of objective and reproducible predictors of AD response could reduce the time needed to achieve remission and relieve patients’ suffering (3). Prior studies suggest that AD response and remission are heritable traits (4), offering the opportunity to use genetic markers to develop predictors applicable in clinical practice to guide drug prescription. The combination of clinical presentation, genomic information and metabolic characteristics was indeed suggested as a possible strategy for the development of precision psychiatry (5).

The purpose of this study was to develop a new approach aiming to contribute to precision psychiatry. Previous studies have focused on the identification of genetic variants associated with AD efficacy (6)(7), and here we expand the focus to transcriptomic profiles derived from transcriptomic-wide association studies (TWAS). Transcriptomic profiles associated with the efficacy of specific ADs in clinical trials can be compared with the *in vitro* AD-induced gene expression changes, in order to test if drug-induced gene expression signatures could be used as markers of clinical efficacy of specific ADs. In this study, we developed and tested this approach by computing gene expression profile associated with remission to citalopram in the STAR*D study and comparing this profile with citalopram and other ADs induced transcriptional responses available from the Connectivity Map (CMap) database. CMap is a genome-scale library of cellular signatures and catalog of transcriptional responses to chemical and genetic perturbations (8). A positive correlation between expression profiles of citalopram remission and *in vitro* citalopram induced gene changes was hypothesized to be indicative of potential utility of our approach. We also hypothesized that the same would be true for escitalopram since it is the therapeutically active enantiomer of citalopram (9).

## Methods

### 1. Study Population

This study is based on Sequence treatment alternative to relieve depression (STAR*D) data (10). The STAR*D study is a trial of protocol-guided antidepressant treatment for outpatients with MDD. The study included 4,041 treatment-seeking adult outpatients, recruited in 18 primary care and 23 psychiatric clinical sites across the United States. Genotyping was performed in 1,948 participants (11). Our analysis used data from the first treatment step (level 1), which consisted of protocol-guided citalopram (20–60 mg/day). Remission was defined as a score < 6 on the Quick Inventory of Depressive Symptomatology clinician-rated (QIDS-C) scale at level 1 exit (after 12 weeks of citalopram treatment). STAR*D genotype and phenotype data are available through the National Institute of Mental Health Human Genetic Initiative (https://www.nimhgenetics.org/). Further details about the STAR*D study are available in the supplementary material (Section 1).

### 2. Genotyping, quality control and imputation

Details on the genotyping procedure can be found elsewhere (11). Individual genotype data was processed using the Psychiatric Genomics Consortium (PGC) “RICOPILI” pipeline for standardized quality control and imputation (12). Imputation of SNPs and insertion-deletion polymorphisms was performed using the 1000 Genomes Project multi-ancestry reference panel (see supplementary material, Section 2).

### 3. Statistical analysis

#### 3.1 Genome wide Association Study (GWAS)

A GWAS was conducted using the RICOPLI pipeline to test the association of each SNP with remission to citalopram, classifying STAR*D participants as remitters or non-remitters. The logistic regression analysis included covariates of sex, age, baseline QIDS-C score and the first 20 population principal components. The GWAS summary statistics were then converted to LD-score regression format using the munge_sumstats.sh script, removing SNPs with an INFO < 0.3 (https://github.com/bulik/ldsc/wiki).

#### 3.2 Transcriptome wide Association Study (TWAS)

We used STAR*D GWAS summary statistics to perform a TWAS using FUSION software (13). Briefly, FUSION requires pre-computed gene expression SNP-weights and GWAS summary statistics to predict the association between the expression of each gene and the phenotype of interest. SNP-weights from CommonMind Consortium dorsolateral prefrontal cortex (DLPFC), 48 tissues within the Genotype-Tissue Expression (GTEx) consortia, Young Finn study, Netherland twin registry, and Metabolic syndrome in men study datasets were considered (Supplementary Table S1). All gene expression SNP-weights were downloaded from the FUSION website (http://gusevlab.org/projects/fusion/). This study uses the term *SNP-weight sets* to define SNP-weights from a given sample and tissue (e.g. GTEx hippocampus, CMC DLPFC). Furthermore, each gene within a given SNP-weight set constitutes a *feature* or *gene-tissue pair.* We combined the FUSION output for all SNP-weight sets, using the TWAS associations (*z*-scores) to represent the gene expression signature of citalopram remitters. The 52 SNP-weight sets in this study contained 252,878 features, representing 26,363 unique genes. Where multiple features for a single gene were available, only the feature providing the highest cross-validation coefficient of determination (CV R^2^) was retained. Similar criteria has been implemented elsewhere (14)

#### 3.3 Comparison of TWAS results with *in vitro* AD-induced gene expression

We evaluated the correlation between the TWAS expression profile of citalopram remission and *in vitro* gene expression profiles of 21 antidepressants available in CMap (Phase II data) (Figure 1-A). CMap is a publicly available comprehensive library of transcriptional expression data obtained using L1000 assay, which directly measures or infers the expression levels of 12,328 genes (https://www.broadinstitute.org/connectivity-map-cmap). The database contains L1000 profiles from various perturbating agents (small molecule compounds, shRNAs, cDNAs, and biologics). More specifically, the CMap platform provides the transcriptomic information of human cultured cell lines exposed to compounds obtained from various screening libraries including drugs approved from FDA (15).

**Figure 1.**
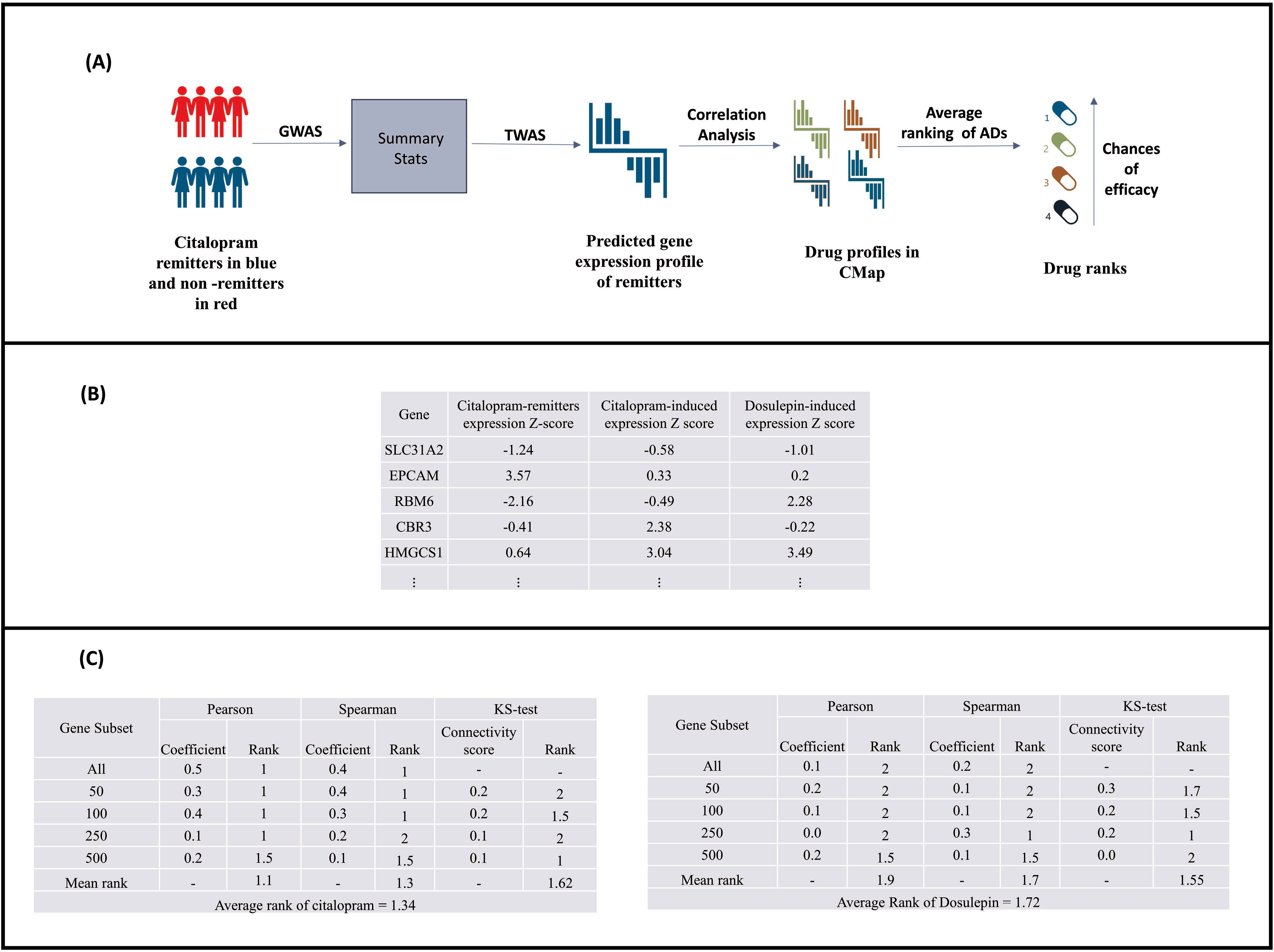
(A) Major steps of the proposed in-silico method. (B) Z-scores of differentially expressed genes of citalopram remitters, dosulepin and citalopram induced profiles from CMap. (C) Description of average rank calculation method using Pearson, Spearman correlation and KS method.

We considered the expression profiles of 21 ADs (Supplementary Table S2) in 5 human cell lines available in Phase II of CMap ((*a) A375, Human malignant melanoma (b) MCF7, Breast cancer (c) PC3, prostate cancer (d) HA1E, kidney (e) HT29, colon cancer).* Drug-induced expression profiles were evaluated in cells treated for 24 hours with 10μm drug concentration. We used CMap’s GEO series (GSE70138) data and extracted relevant expression profiles using cmapR package. Of the 12,328 genes within the CMap profiles, 10,027 were captured by the SNP-weight included in the citalopram remitter TWAS. We compared the expression profiles of the 21 ADs with the profile of citalopram remitters obtained from the TWAS using an approach described in a previous study (16). The differentially expressed genes represented in terms of z-scores of citalopram remitters and drug induced profiles (Figure 1-B) were analyzed using R code (https://sites.google.com/site/honcheongso/software/gwascmap), according to the following procedure:

a. *Evaluating the relationship between AD-induced gene expression and expression profiles of citalopram remitters*. Patterns of expressions were tested by analyzing all and the strongest up regulated and down regulated genes in the TWAS (*k* = 50, 100, 250, 500). The correlation between CMap antidepressant profiles and the STAR*D remitter profile was assessed for each drug using Spearman’s correlation and Pearson’s correlation using all and highly modulated remitter’s *k* genes. We adopted the KS test as reported by the original CMap study to compare the expression patterns of AD and citalopram remitters by considering strongly up and downregulated *k* genes and calculated connectivity scores (8). The 21 tested ADs were ranked based on the results of each test (Pearson, Spearman and KS), and then the average rank across tests for each drug were computed. Drugs were ranked in ascending order of their correlation results (the drug with most positive correlation was ranked first) (Figure 1-C).
b. *Significance of ranks using permutation.* In order to estimate the significance of the ranks, a permutation procedure was performed by shuffling the z-scores obtained in the TWAS and calculating the corresponding rank of each drug by repeating the procedure in step a. One hundred permutations were performed to calculate the distribution of ranks under the null hypothesis and estimated the p-value of the observed ranks.
c. *Calculation of ranks probability for each AD across cell lines using Genome Scan Meta-Analysis (GSMA) method*. We combined ranks of each AD in five cell lines by adding them and calculated the sum of ranks probability using GSMA, a non-parametric method for meta-analyzing ranks (17)

Finally, we repeated the process described in a. and b. for five control drugs (Supplementary Table S3) having hypothetically no antidepressant effect in order to validate the proposed method. The major steps of the applied in-silico method are shown in Figure 1-A.

## Results

STAR*D data included 506 citalopram remitters and 657 non-remitters with genotypic data after quality control, and the main clinical-demographic characteristics are shown in (Supplementary Table S4). The GWAS and TWAS Q-Q plots showed no evidence of confounding. (Supplementary Figures S1 and S2).

The average rank across tests for ADs showed that escitalopram (S-enantiomer of citalopram) was the AD with the highest average rank followed by amitriptyline in MCF7 (breast cancer cell line). In A375 (human malignant melanoma) and PC3 (prostate cancer) cell lines, escitalopram and citalopram emerged as the second highest ranked ADs, respectively, after trimipramine and mirtazapine, respectively. In HT29 (colon cancer) cell line, citalopram ranked third after trimipramine and dosulepin. Imipramine and fluvoxamine were the top ranked ADs in HA1E (kidney) cell line, whereas escitalopram and citalopram did not fall in the top ranks in this cell type. In the analysis of combined ranks across cell lines, we found sertraline, trimipramine and venlafaxine as drugs with the best sum of ranks and *p-values* < 0.05, while citalopram was right after them and close to the significance threshold (*p=0.057*) (Table 1).

**Table 1.** Ranking of ADs in five human cell lines. P-values were obtained through the permutation procedure described in section 3.3. P values < 0.05 are highlighted in each cell line and in the combined cell line results.

| Cell lines | A375 |  | MCF7 |  | PC3 |  | HA1E |  | HT29 |  | Combined cell lines |  |
| --- | --- | --- | --- | --- | --- | --- | --- | --- | --- | --- | --- | --- |
| Drug | Rank | Perm.p.value | Rank | Perm.p.value | Rank | Perm.p.value | Rank | Perm.p.value | Rank | Perm.p.value | Rank Sum | p.value |
| Amitriptyline | 16.9 | 0.8610 | 2.4 | 0.0290 | 12.6 | 0.6138 | 17.6 | 0.8948 | 12.9 | 0.6224 | 62.4 | 0.705 |
| Citalopram | 7.6 | 0.2862 | 7.7 | 0.2890 | 2.3 | 0.0276 | 10.6 | 0.4776 | 4.4 | 0.1114 | 32.6 | 0.057 |
| Clomipramine | 11.1 | 0.5229 | 4.1 | 0.0890 | 9.1 | 0.3714 | 18.6 | 0.9371 | 16.8 | 0.8590 | 59.7 | 0.653 |
| Dosulepin | 17.3 | 0.8824 | 18.1 | 0.9257 | 15 | 0.7605 | 8.8 | 0.3571 | 2.6 | 0.0405 | 61.8 | 0.705 |
| Duloxetine | 11 | 0.5181 | 20.9 | 0.9971 | 17.2 | 0.8776 | 11.3 | 0.5305 | 4.8 | 0.1300 | 65.2 | 0.775 |
| Escitalopram | 2.6 | 0.0310 | 1.2 | 0.0014 | 6.9 | 0.2395 | 14.5 | 0.7324 | 17.6 | 0.8976 | 42.8 | 0.203 |
| Fluoxetine | 13.8 | 0.6843 | 10.8 | 0.4890 | 17.8 | 0.9033 | 6.2 | 0.1976 | 16.7 | 0.8562 | 65.3 | 0.775 |
| Fluvoxamine | 19.6 | 0.9729 | 6.2 | 0.2010 | 2.9 | 0.0443 | 3.2 | 0.0529 | 9.6 | 0.4005 | 41.5 | 0.183 |
| Imipramine | 6.2 | 0.2014 | 12.6 | 0.6052 | 14.6 | 0.7395 | 1.2 | 0.0062 | 5.2 | 0.1490 | 39.8 | 0.147 |
| Maprotiline | 3.7 | 0.0667 | 6.8 | 0.2362 | 9.8 | 0.4200 | 11.2 | 0.5229 | 10.5 | 0.4590 | 42 | 0.183 |
| Mianserin | 19.6 | 0.9729 | 15.6 | 0.8029 | 17.8 | 0.9033 | 17.9 | 0.9062 | 19.9 | 0.9795 | 90.8 | 0.998 |
| Mirtazapine | 9.8 | 0.4362 | 18.8 | 0.9529 | 2 | 0.0181 | 7.7 | 0.2871 | 5.2 | 0.1490 | 43.5 | 0.225 |
| Nortriptyline | 11.6 | 0.5576 | 15.2 | 0.7833 | 12 | 0.5748 | 18.1 | 0.9167 | 7.9 | 0.2990 | 64.8 | 0.775 |
| Paroxetine | 18.3 | 0.9248 | 14.4 | 0.7390 | 6.4 | 0.2129 | 14.7 | 0.7448 | 16.7 | 0.8562 | 70.5 | 0.870 |
| Reboxetine | 11.8 | 0.5681 | 10.2 | 0.4476 | 10.7 | 0.4895 | 19.8 | 0.9800 | 13.3 | 0.6552 | 65.8 | 0.797 |
| Selegiline | 10.6 | 0.4890 | 17 | 0.8667 | 19.2 | 0.9581 | 15 | 0.7671 | 15.6 | 0.8005 | 77.4 | 0.951 |
| Sertraline | 4.4 | 0.1033 | 6.3 | 0.2052 | 9.1 | 0.3714 | 3.6 | 0.0662 | 7.4 | 0.2743 | 30.8 | 0.041 |
| Tranlycypromine | 16.9 | 0.8610 | 14.8 | 0.7619 | 18 | 0.9119 | 9.5 | 0.3929 | 16.9 | 0.8614 | 76.1 | 0.943 |
| Trazodone | 10.3 | 0.4657 | 10.9 | 0.4943 | 13 | 0.6424 | 8 | 0.3043 | 16.6 | 0.8529 | 58.8 | 0.626 |
| Trimipramine | 2.4 | 0.0267 | 13.4 | 0.6614 | 4 | 0.0919 | 9.7 | 0.4086 | 1.4 | 0.0105 | 30.9 | 0.041 |
| Venlafaxine | 5.5 | 0.1567 | 3.6 | 0.0676 | 10.6 | 0.4819 | 3.8 | 0.0810 | 9 | 0.3657 | 32.5 | 0.049 |

We also attempted to validate our approach using the expression profiles of five control drugs. In A375 all control drugs ranked after (es)citalopram. In PC3, three control drugs ranked after (es)citalopram. In MCF7, four control drugs were ranked after escitalopram and one control drug was listed after citalopram. These observations support the hypothesis that expression profiles associated with remission to a specific AD are on average more correlated with *in vitro* gene expression induced by the same AD than that induced by other drugs. In HT29 and HA1E, four control agents ranked before the (es)citalopram. The rankings of ADs together with control drugs in the five analyzed cell lines are showed in (Supplementary Tables S5-9).

We observed that AD-induced expression profiles vary across the five analyzed cell lines. Interestingly, citalopram and escitalopram have distinctive signatures, with a weak correlation between them in four cell lines (A375, MCF7, PC3 and HT29), and moderate correlation in one cell line (r=0.200 for HA1E) (Table. 2). The observed variability in drug-induced gene expression among cell lines likely contributes to the differences in ADs ranking across cell lines. The variability of all AD-induced profiles across cell lines are reported as correlation matrices in the supplementary file (Supplementary Figures S3-12).

**Table 2.** Spearman correlation between citalopram and escitalopram in each cell line. P-value suggesting correlation coefficient being different from zero.

| Cell lines | Correlation coefficient | p-value |
| --- | --- | --- |
| A375 | -0.008 | 0.405 |
| MCF7 | 0.038 | 9.74E-05 |
| PC3 | -0.03 | 0.002 |
| HT29 | -0.01 | 0.298 |
| HA1E | 0.2 | 1.34E-91 |

## Discussion

AD response is heterogeneous among MDD patients and no objective and validated predictors of response are yet available (18). In this study, we evaluated an in-silico approach to prioritize ADs utilizing imputed gene expression profiles of citalopram remitters and AD-induced transcriptional profiles available in CMap. The positive correlation between gene expression of citalopram remitters and (es)citalopram induced expression profiles in three cell lines (A375, MCF7 and PC3) suggests that the predicted gene expression profile of a remitter is correlated with *in vitro* expression profiles induced by the same ADs. No previous study has tested this hypothesis and our results show that approach might be used to rank ADs based on their likelihood of efficacy for an individual.

Analysis of transcriptional profiles of drugs and diseases signatures is already an established approach in the domain of drug repositioning. For instance, Sirota and colleagues found that cimetidine showed an opposite expression pattern to that associated with lung adenocarcinoma and experimentally validated this drug as a potential treatment (19). Similarly, topiramate was found as a possible treatment for inflammatory bowel disease and this hypothesis was validated in an animal model (20). In our study, we applied a similar strategy, but instead of disease-associated gene expression signatures we used expression profiles associated with remission to a known AD drug to test if they could be useful to prioritize the prescription of available ADs, rather than for identifying new potential ADs. We indeed hypothesized that AD induced expression profiles *in vitro* may be correlated with gene expression profiles observed in remitters to the same drug and similar drugs.

A major observation that emerged from our results was the difference in the ranking of ADs drugs across cell lines which suggested that the selection of the most appropriate cell line(s) is a relevant step for applying our approach. The differences in results among cell lines can be explained by the inter-cellular drug induced expression signature variability, as reported by Subramanian and colleagues. According to this study, 15% of all the drug compounds produced highly similar signatures across 9 cell lines, whereas the remaining drugs produced diverse signatures (15). The heterogeneity of drug signatures depends on the cellular pathways associated with a cell type. In this study we observed that A375 and MCF7 provided results which were more consistent with our hypothesis compared with other cell lines for both ADs and control drugs. This may be explained by the similar embryological origin of these cell lines. A375 and MCF7 are skin and breast cancer cell lines respectively, and both skin and breast cells originate from the ectoderm (outermost layer of embryo), the same layer from which nervous tissues originate (21). This hypothesis suggests that the use of brain cell lines may be more suitable for our study, but this was not possible as discussed among the limitations.

Despite the low comparability of gene expression profiles across cell lines, we decided to calculate the significance of cumulative ranks across cell lines. By combining ranks of ADs in the evaluated cell lines, the evidence of association of remitters’ signature to drugs other than (es)citalopram might indicate that they may induce a similar gene expression profile to that observed in remitters to citalopram, thus hypothetically patients who benefit from citalopram may benefit also from these ADs.

Citalopram is a racemic mixture comprised of two enantiomers, R and S-citalopram (escitalopram) in equal proportions. However, the signatures of citalopram and escitalopram are only weakly correlated in the five analyzed cell lines. This can be due to the differences in modulated genes and pathways by these drugs *in vitro*, as reported by Sakka et al. Their study suggests that citalopram and escitalopram modulated 69 and 42 pathways, respectively, and 10 pathways were differentially modulated in a neuroblastoma cell line (22). In other words, the *in vitro* gene expression profile of citalopram is influenced by both escitalopram and R-citaloram to a similar extent, making it different compared to the profile of escitalopram alone. On the other hand, the *in vivo* gene expression signature of citalopram remitters is hypothetically highly dependent on genes regulated by escitalopram rather than R-citalopram, since escitalopram has a 50-fold higher affinity for the serotonin transporter compared to R-citalopram and it is considered responsible for the therapeutic effects of citalopram (23). However, escitalopram was not close to significance in the analyses of combined ranks across cell lines.

Our approach is innovative and shows important strengths. First, it reflects the polygenic architecture of AD response, characterized by multiple effects of small size (7). Second, it is based on genotype-predicted gene expression profiles providing an advantage over traditional expression data from microarray and RNA sequencing methods. Patients from expression studies are indeed mostly medicated and brain tissues can only be acquired from postmortem samples, hence, psychiatric medications might confound the expression results. On the contrary, genotype-predicted gene expression profiles are not susceptible to alteration due to medications because this approach only captures the heritable component of gene expression. Additionally, GWAS sample sizes are usually significantly larger than those used in expression studies and GWAS summary statistics are easily accessible for a number of traits. Further, the expression profiles can be imputed for different tissues which can help to comprehend biological mechanisms at the tissue level. Last, our method is computationally simple, and it can be applied to other traits.

There are also some limitations of the proposed methodology. First, this method can be implemented only at population level (on data from an aggregated sample of individuals), while it needs to be adapted for application at the individual level. Second, we could not test our method in neuronal progenitor cells or differentiated neurons from the CMap transcriptional catalogue since AD-induced gene expression for these cell lines was not available. A prior CMap study suggested that neuronal cell lines are different compared to cancer cell lines in terms of the drug expression profiles but neuropsychiatric diseases can be reasonably modeled using cancer cell lines (15)(8). However, the relevant differences between expression profiles of different cell lines found by our study and previous studies suggest that the identification of the most suitable cell line for the trait of interest is an important step. Neural cell lines may indeed show distinctive pathways which are relevant to AD action.

In conclusion, we tested an in-silico approach in five human cell lines by using GWAS results and drug-induced profiles to rank ADs based on their correlation, which hypothetically may reflect the chances of efficacy of specific ADs. This study indicates that there is a correlation between (es)citalopram induced expression profiles and predicted expression associated with remission to citalopram only in some cell lines. Therefore, at the individual level, on average the predicted expression of (es)citalopram remitters should be more correlated with (es)citalopram-induced expression than non-remitters. Our approach can further be extended by investigating the correlation between a drug-induced expression profile and an individual’s predicted gene expression levels which can be used to rank drugs by their predicted efficacy. Hence, the given method can be improved by considering genotype data at the individual level and using expression signatures of brain cell lines. A ‘one size fits all’ is not a valid strategy for the treatment of MDD and our study proposed a new approach to contribute to the development of precision psychiatry.

## Acknowledgements

We thank the NIMH for providing the opportunity of analyzing their data on the STAR*D sample. We would like to thank Connectivity map team for making their data available for community research use. This paper represents independent research part-funded by the National Institute for Health Research (NIHR) Biomedical Research Centre at Oxford, South London, Maudsley NHS Foundation Trust and King’s College London. The views expressed are those of the authors and not necessarily those of the NHS, the NIHR or the Department of Health and Social Care. We acknowledge the use of research computing facility at King’s College London, Rosalind (https://rosalind.kcl.ac.uk), which is delivered in partnership with the National Institute for Health Research (NIHR) Biomedical Research Centre at South London & Maudsley and Guy’s & St. Thomas’ NHS Foundation Trusts, and part-funded by capital equipment grants from the Maudsley Charity (award 980) and Guy’s & St. Thomas’ Charity (TR130505).

## Supplementary information

Supplementary information is available at The Pharmacogenomics Journal’s website

