## Supplementary file for "Investigating an in-silico approach for prioritizing antidepressant drug prescription based on drug-induced expression profiles and predicted gene expression"

### Supplementary figures list

Figure S1. GWAS QQ- plot for citalopram remission

Figure S2. TWAS QQ- plot for citalopram remission

Figure S3. Correlation matrix plot between AD signatures of A375 and MCF7

Figure S4. Correlation matrix plot between AD signatures of A375 and PC3

Figure S5. Correlation matrix plot between AD signatures of A375 and HT29

Figure S6. Correlation matrix plot between AD signatures of A375 and HA1E

Figure S7. Correlation matrix plot between AD signatures of MCF7 and HT29

Figure S8. Correlation matrix plot between AD signatures of MCF7 and PC3

Figure S9. Correlation matrix plot between AD signatures of MCF7 and HA1E

Figure S10. Correlation matrix plot between AD signatures of HA1E and PC3

Figure S11. Correlation matrix plot between AD signatures of HT29 and PC3

Figure S12. Correlation matrix plot between AD signatures of HA1E and HT29

**Supplementary tables list**

Table S1.  Tissues considered for TWAS analysis.

Table S2. Antidepressants and drug classes

Table S3. Control agents and drug classes

Table S4. Main clinical demographic characteristics of STAR*D

Table S5. Ranking of ADs and control drugs in A375

Table S6. Ranking of ADs and control drugs in MCF7

Table S7. Ranking of ADs and control drugs in PC3

Table S8. Ranking of ADs and control drugs in HA1E

Table S9. Ranking of ADs and control drugs in HT2

**Supplementary material**

*Section 1: Cohort information*

Sequence treatment alternative to relieve depression (STAR*D) was a collaborative study supported by National institute of mental health to study different treatment strategies in real world MDD patients. The STAR*D study recruited patients between the age of 18-75 from psychiatric and primary health care clinics. The trial continues for about four years, started in 2000 with the enrollment of patients suffering from non-psychotic depressive disorder and completed with their follow-up in 2004 (1). The study design of STAR*D comprised of four treatment levels to assess treatment response. The time period for each level was 14 weeks. The total enrolled 4,000 individuals started from level 1, if the patients didn't achieve significant remission by the end of 14th week of each level, they entered the subsequent stage of treatment (2) (3). Alternatively, patients with symptomatic improvement and remission were excluded from the study and encouraged for the one-year follow-up. Genetic material was collected from 1,948 (48%) participants; of whom 1,491 (37% of the original STAR*D sample, including 980 of white/European ancestry) passed quality control and were included in previously reported genome-wide analyses (4). The study was approved by institutional ethics review boards at all centres. Written consent was obtained from all participants after the procedures and any associated risks were explained.

*Section 2: Quality control and imputation of genotype data using RICOPILI*

Individual genotype data for all cohorts were processed using the PGC "RICOPILI" pipeline for standardized quality control, imputation, and association analysis (5). Quality control and imputation were performed according to the standards from the Psychiatric Genomics Consortium (PGC). The default parameters for retaining SNPs and subjects were: SNP missingness < 0.05 (before sample removal); subject missingness < 0.02; autosomal heterozygosity deviation ($|F_{het}|<0.2$); SNP missingness < 0.02 (after sample removal); difference in SNP missingness between cases and controls < 0.02; and SNP Hardy-Weinberg equilibrium ($P>10^{-6}$ in controls or $P>10^{-10}$ in cases). These default parameters sufficiently controlled $\lambda$ and false positive findings.

Genotype imputation was performed using the pre-phasing/imputation stepwise approach implemented in IMPUTE2 / SHAPEIT (chunk size of 3 Mb and default parameters). The imputation reference set consisted of 2,186 phased haplotypes from the 1000 Genomes Project dataset (August 2012, 30,069,288 variants, release "v3.macGT1"). After imputation, we identified SNPs with very high imputation quality (INFO >0.8) and low missingness (<1%) for building the principal components to be used as covariates in final association analysis. SNPs underwent linkage disequilibrium-based pruning ($r^2 > 0.02$) and frequency filtering (MAF > 0.05). This SNP set was used for robust relatedness testing and population structure analysis. Relatedness testing identified pairs of subjects with $\hat{\pi} > 0.2$, and one member of each pair was removed at random after preferentially retaining cases over controls. Principal component estimation used the same collection of autosomal SNPs.

Identification of identical samples is easily accomplished given direct access to individual genotypes. One concern is the inclusion of closely related individuals. We used SNPs directly genotyped on all platforms to compute empirical relatedness and excluded one of each duplicated or relative pair (defined as $\hat{\pi} > 0.2$).

**Supplementary figures**

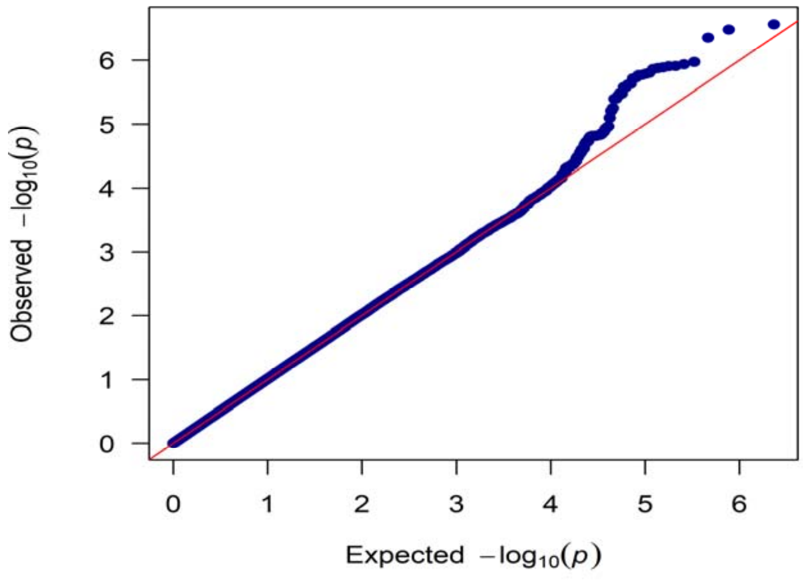

Figure S1. QQ plot of GWAS p-values, N (p-values) = 1158655

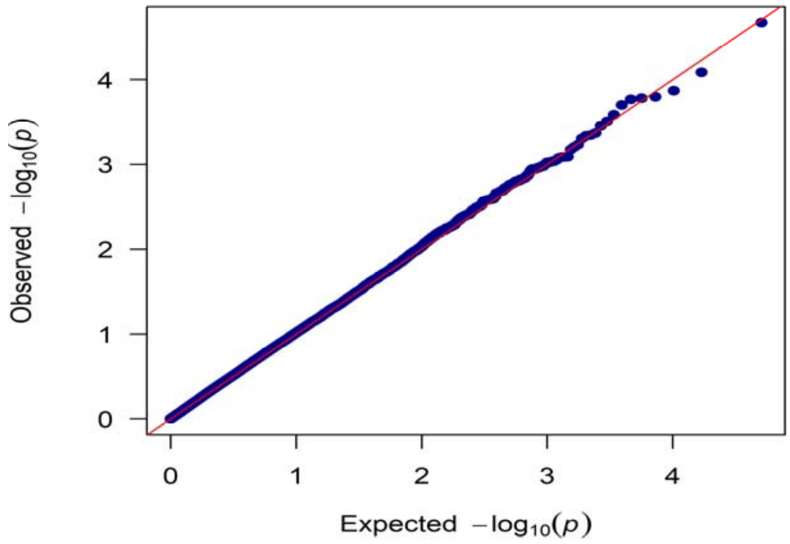

Figure S2. QQ plot of TWAS p-values, N (p-values) = 26363

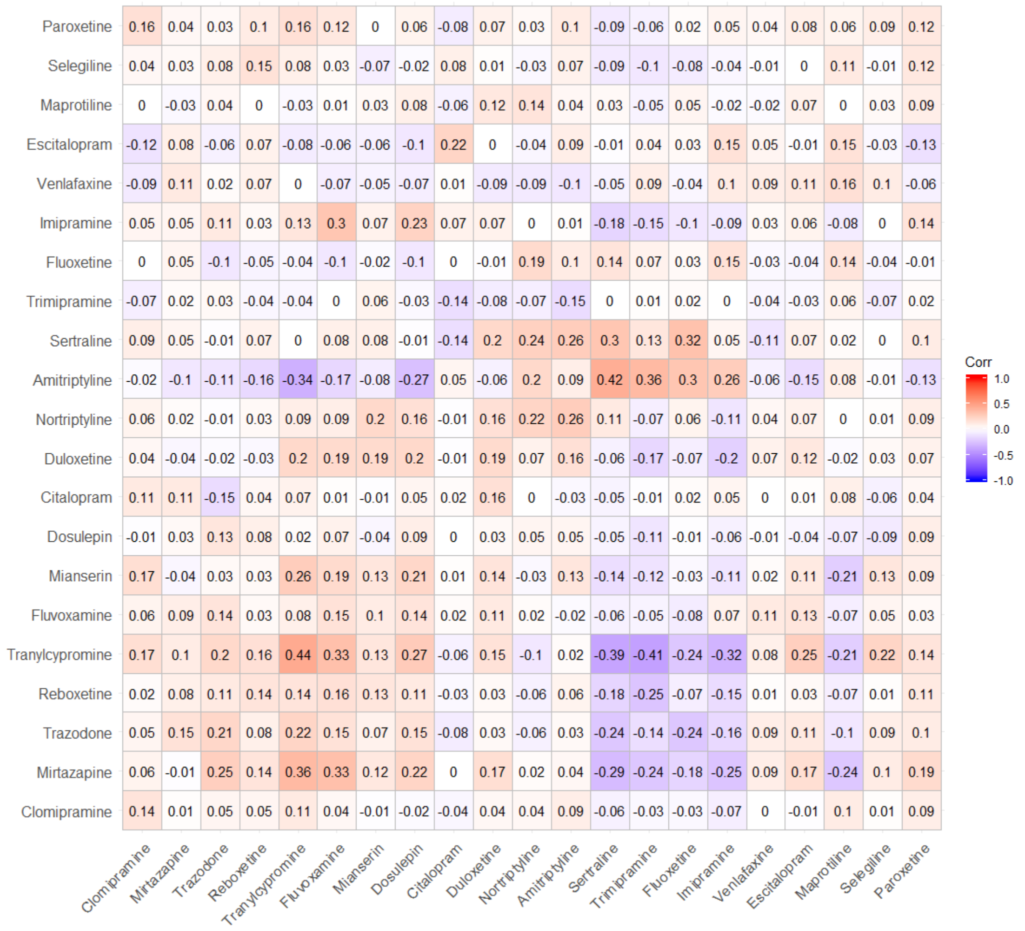

Figure S3. Correlation Matrix plot between AD signatures of A375 and MCF7

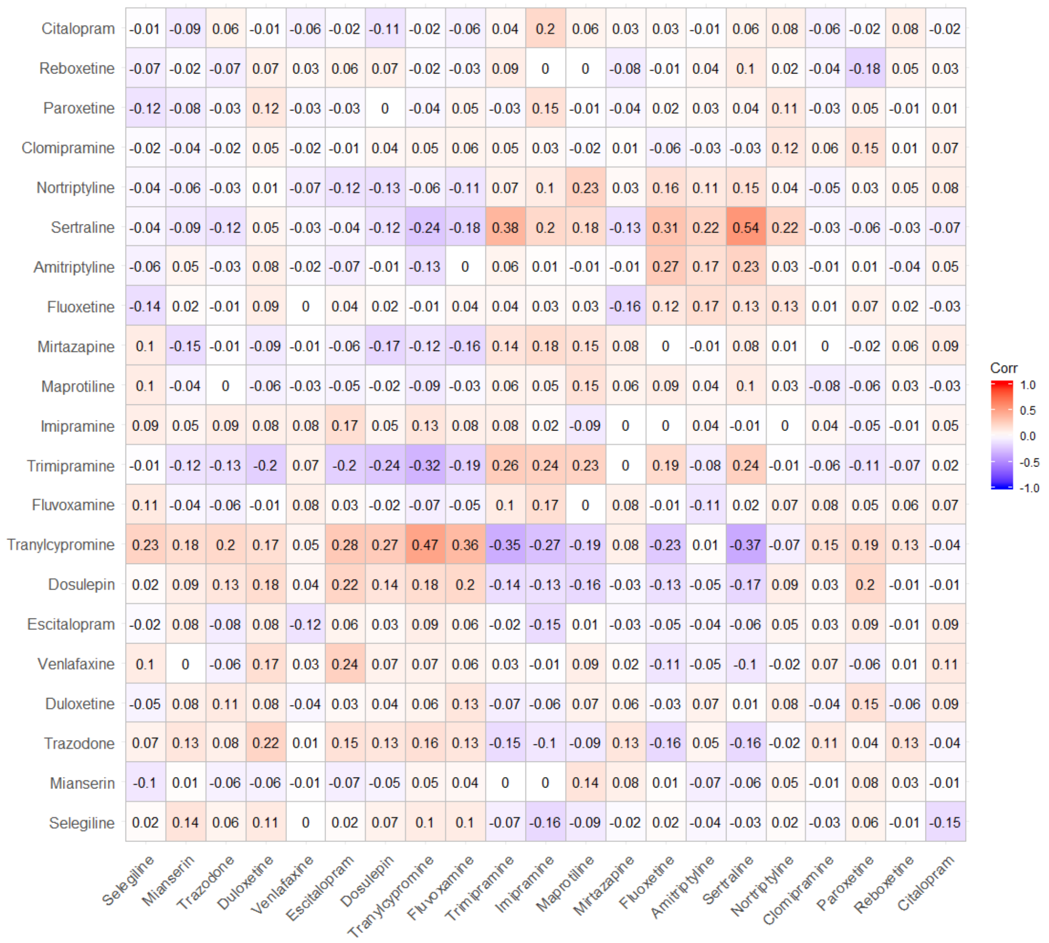

Figure S4. Correlation matrix plot between AD signatures of A375 and PC3

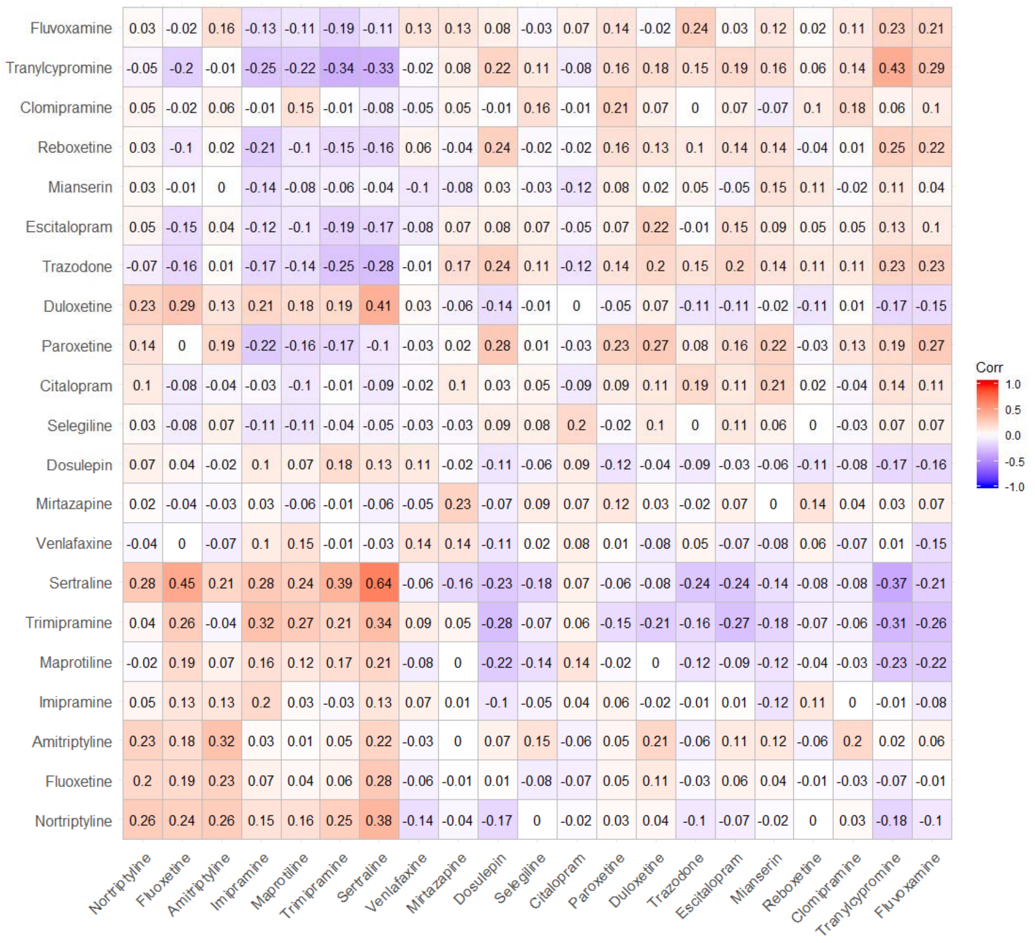

Figure S5. Correlation matrix plot between AD signatures of A375 and HT29

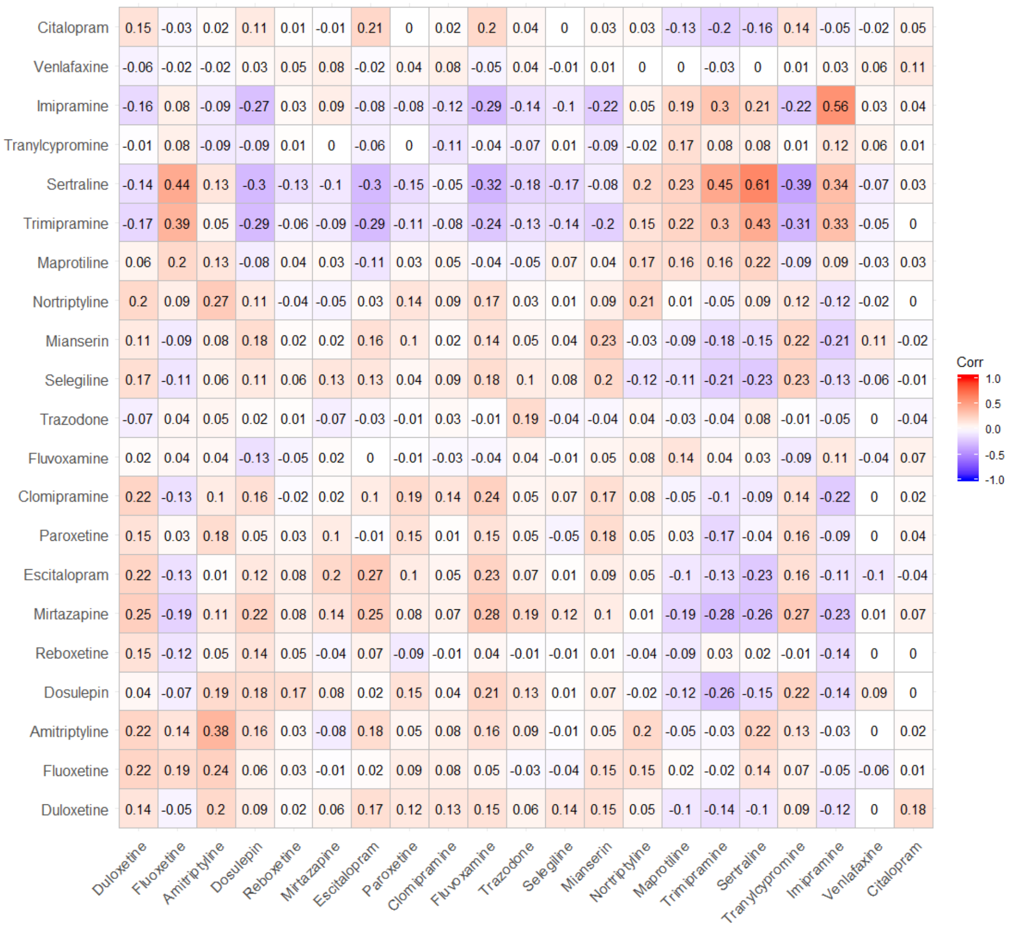

Figure S6. Correlation matrix plot between AD signatures of A375 and HA1E

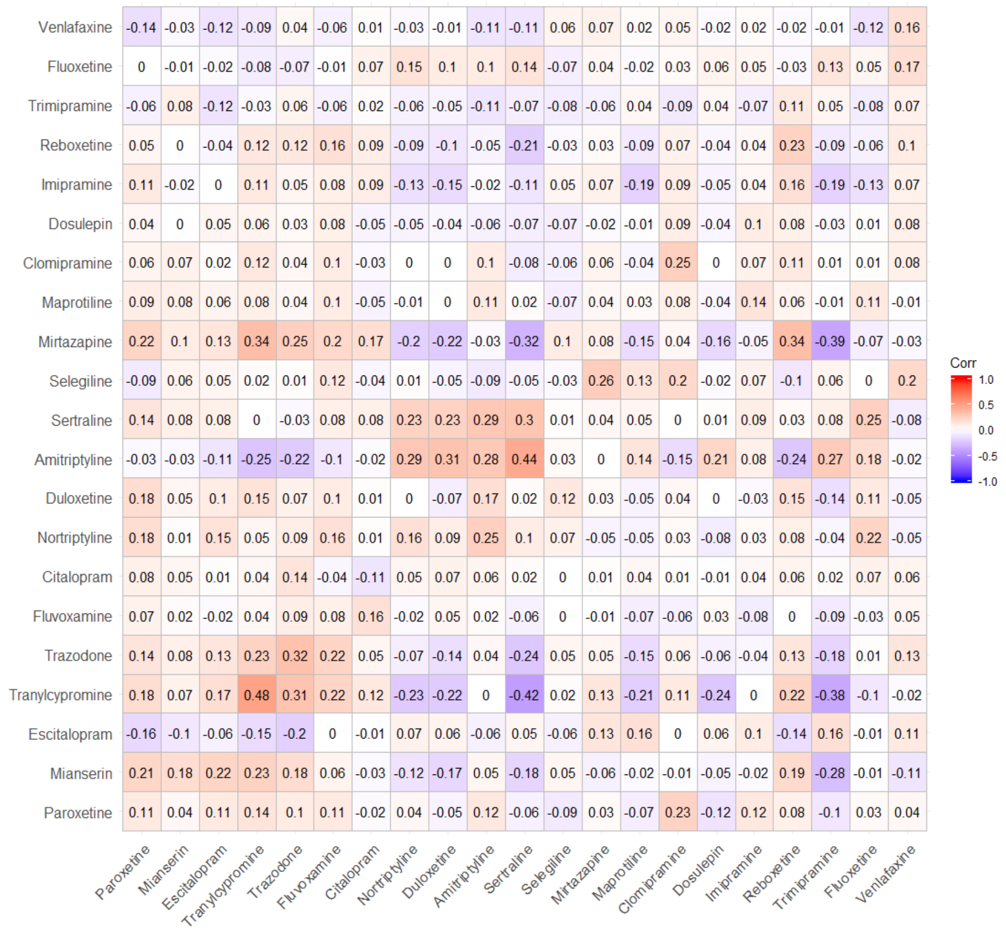

Figure S7. Correlation matrix plot between AD signatures of MCF7 and HT29

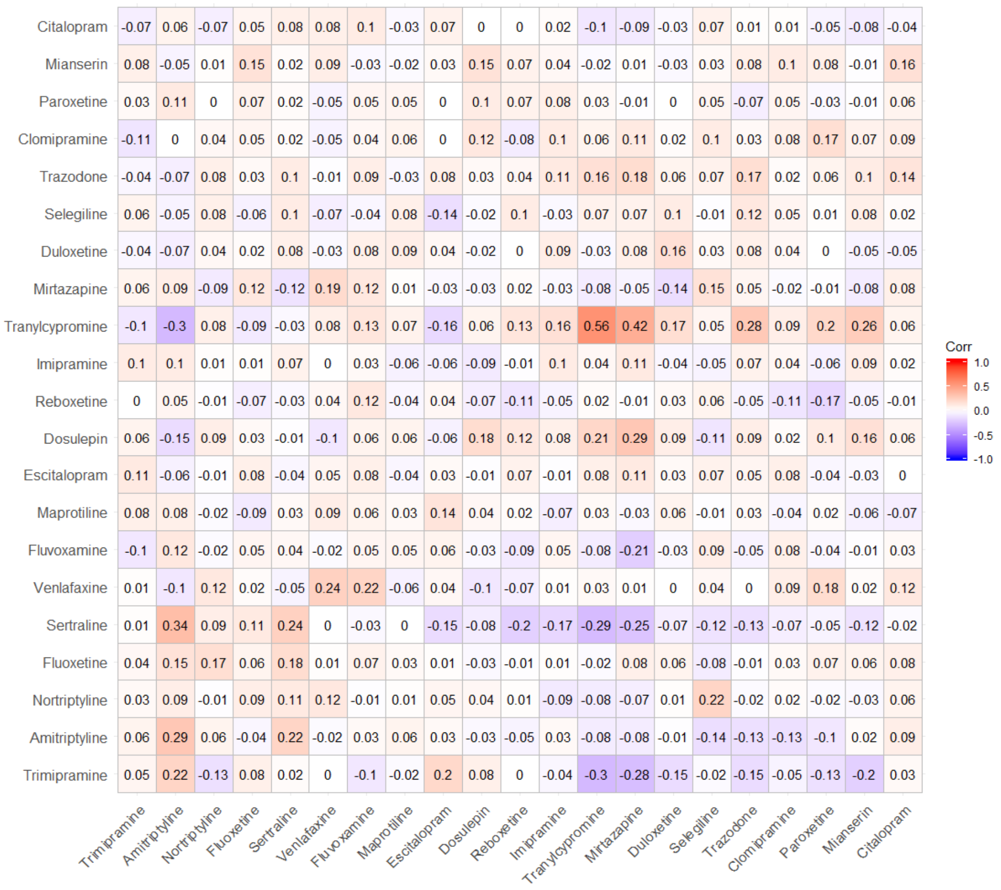

Figure S8. Correlation matrix plot between AD signatures of MCF7 and PC3

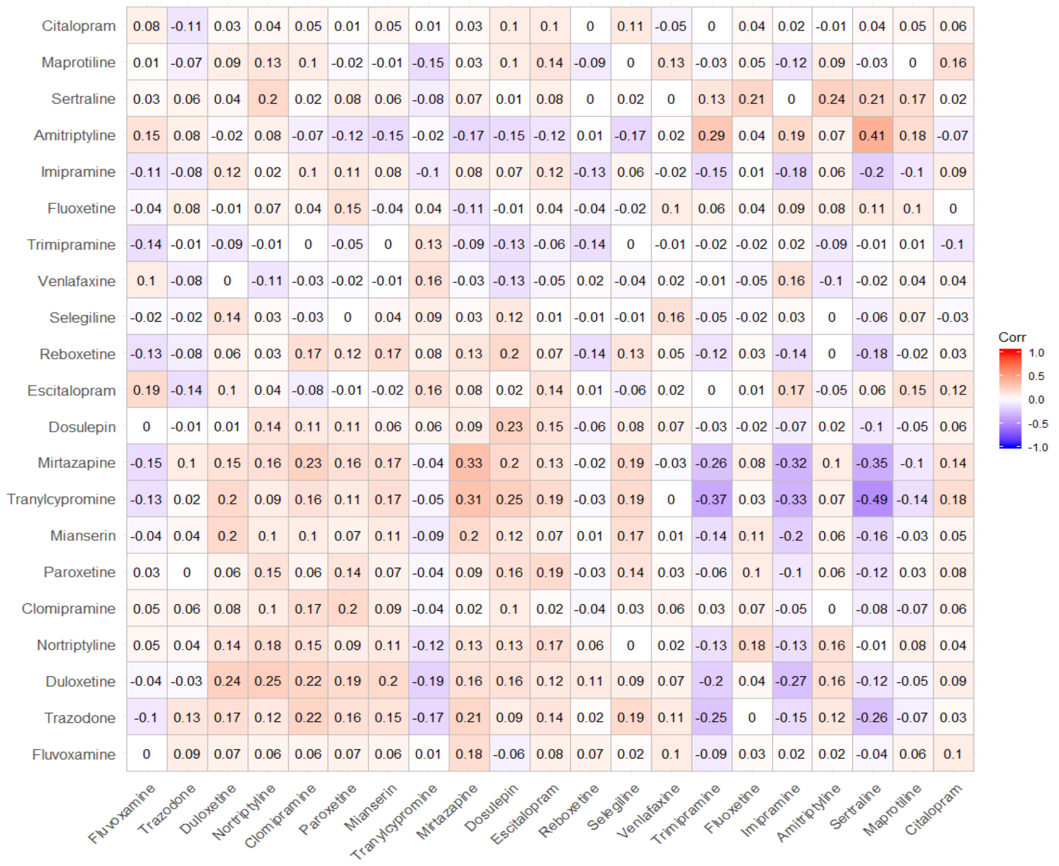

Figure S9. Correlation matrix plot between AD signatures of MCF7 and HA1E

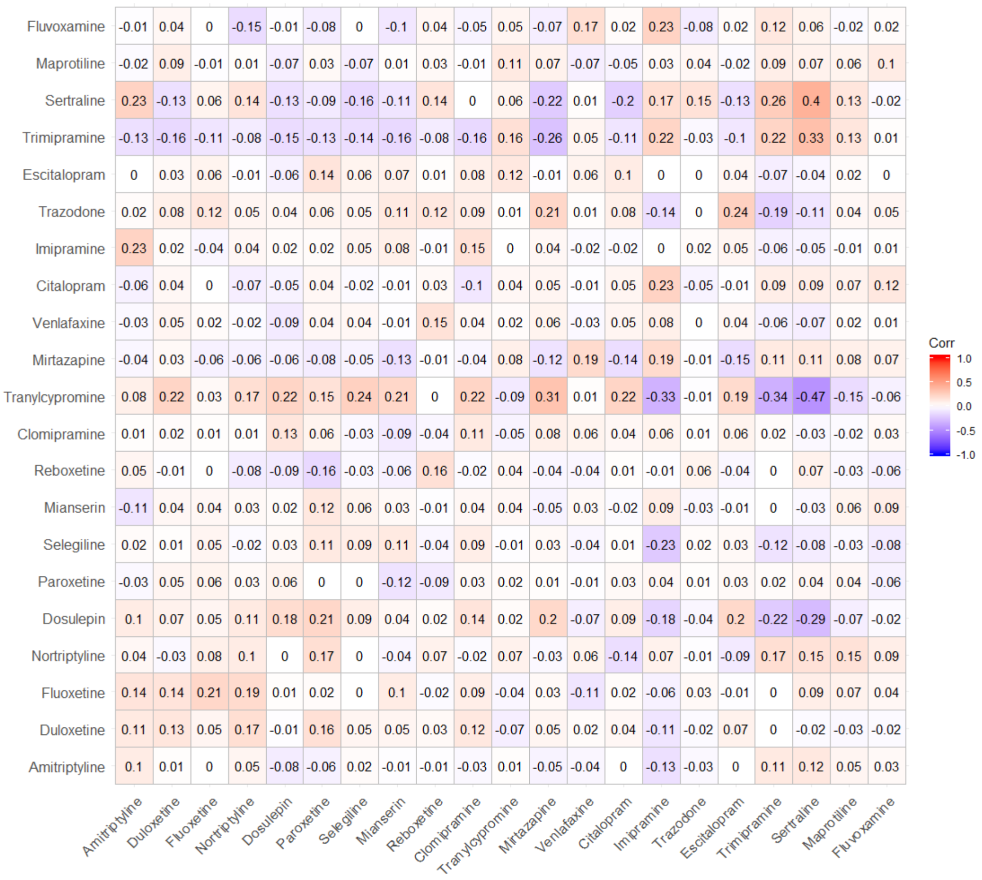

Figure S10. Correlation matrix plot between AD signatures of HA1E and PC3

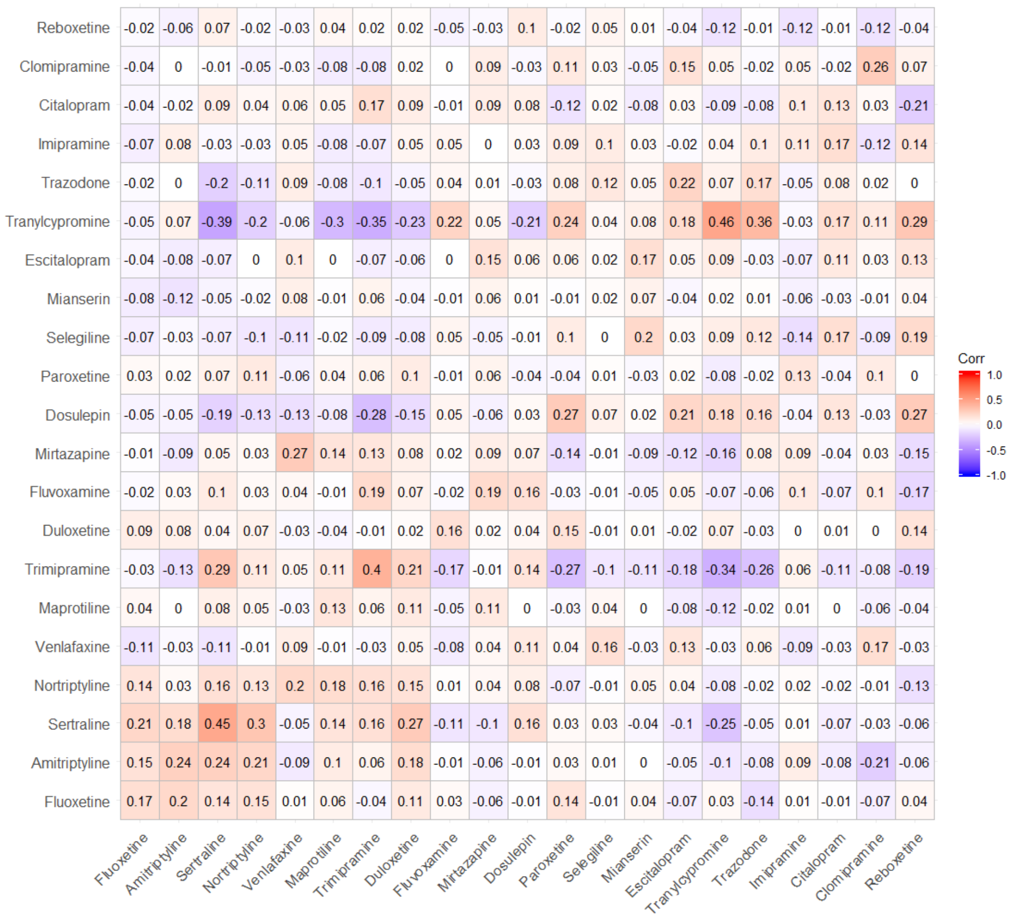

Figure S11. Correlation matrix plot between AD signatures of HT29 and PC3

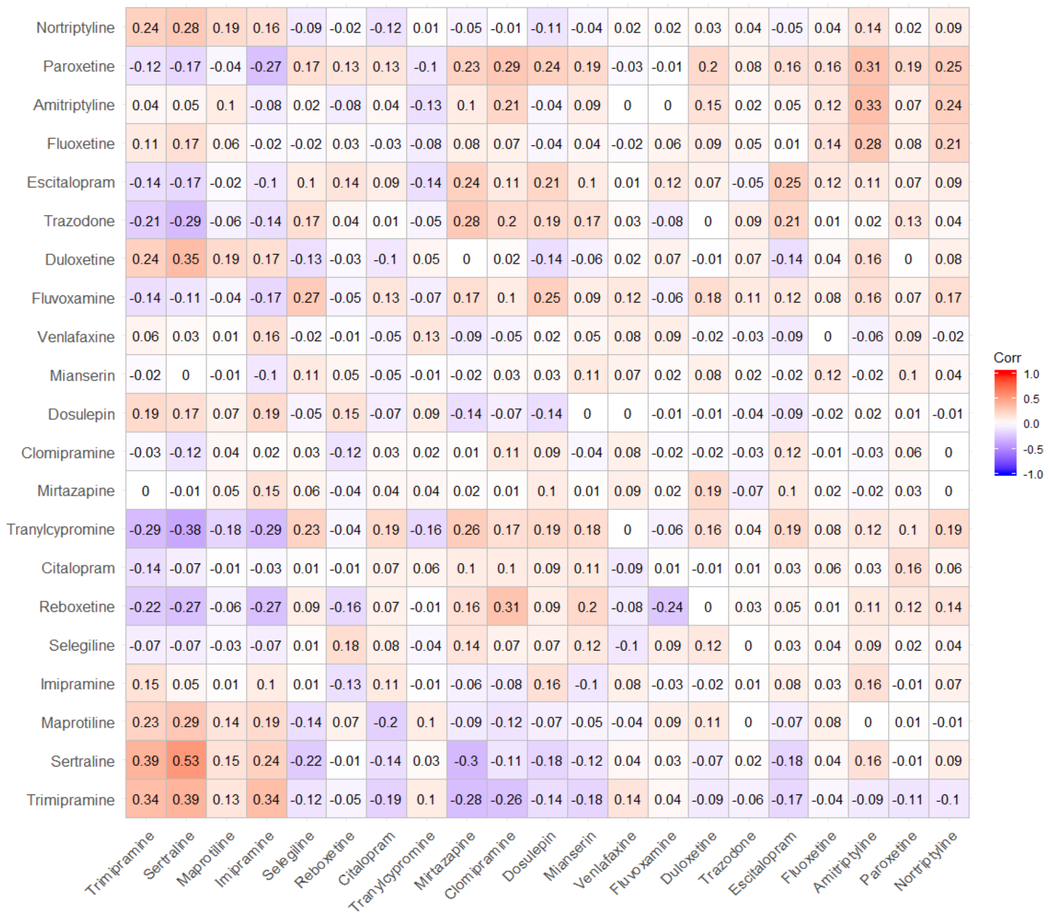

Figure S12. Correlation matrix plot between AD signatures of HT29 and HA1E

### Supplementary Tables

Table S1. Tissues considered for TWAS analysis.

| **GTEx v7 multi-tissue (RNA-seq)** | |
|---|---|
| **Tissue** | **No of Samples** |
| Adipose - Subcutaneous | 385 |
| Adipose - Visceral (Omentum) | 313 |
| Adrenal Gland | 175 |
| Artery - Aorta | 267 |
| Artery - Coronary | 152 |
| Artery - Tibial | 388 |
| Brain - Amygdala | 88 |
| Brain - Anterior cingulate cortex (BA24) | 109 |
| Brain - Caudate (basal ganglia) | 144 |
| Brain - Cerebellar Hemisphere | 125 |
| Brain - Cerebellum | 154 |
| Brain - Cortex | 136 |
| Brain - Frontal Cortex (BA9) | 118 |
| Brain - Hippocampus | 111 |
| Brain - Hypothalamus | 108 |
| Brain - Nucleus accumbens (basal ganglia) | 130 |
| Brain - Putamen (basal ganglia) | 111 |
| Brain - Spinal cord (cervical c-1) | 83 |
| Brain - Substantia nigra | 80 |
| Breast - Mammary Tissue | 251 |
| Blood - EBV-transformed lymphocytes | 117 |
| Skin - Transformed fibroblasts | 300 |
| Colon - Sigmoid | 203 |
| Colon - Transverse | 246 |
| Esophagus - Gastroesophageal Junction | 213 |
| Esophagus - Mucosa | 358 |
| Esophagus - Muscularis | 335 |
| Heart - Atrial Appendage | 264 |
| Heart - Left Ventricle | 272 |
| Liver | 153 |
| Lung | 383 |
| Minor Salivary Gland | 85 |
| Muscle - Skeletal | 491 |
| Nerve - Tibial | 361 |
| Ovary | 122 |
| Pancreas | 220 |
| Pituitary | 157 |

| | |
|---|---|
| Prostate | 132 |
| Skin - Not Sun Exposed (Suprapubic) | 335 |
| Skin - Sun Exposed (Lower leg) | 414 |
| Small Intestine - Terminal Ileum | 122 |
| Spleen | 146 |
| Stomach | 237 |
| Testis | 225 |
| Thyroid | 399 |
| Uterus | 101 |
| Vagina | 106 |
| Whole Blood | 369 |
| **Common mind consortium (RNA seq)** | |
| Brain prefrontal cortex | 452 |
| **Metabolic Syndrome in men (RNA seq)** | |
| Adipose | 563 |
| **Young Finns Study (Expression microarray)** | |
| Blood | 1264 |
| **Netherland twin registry (Expression microarray)** | |
| Blood | 1247 |

Table S2. List of Antidepressants and drug class

| Antidepressants | Drug Class |
|---|---|
| Citalopram | Selective serotonin reuptake inhibitor |
| Escitalopram | Selective serotonin reuptake inhibitor |
| Fluoxetine | Selective serotonin reuptake inhibitor |
| Fluvoxamine | Selective serotonin reuptake inhibitor |
| Paroxetine | Selective serotonin reuptake inhibitor |
| Sertraline | Selective serotonin reuptake inhibitor |
| Trazodone | Serotonin antagonist and reuptake inhibitor |
| Duloxetine | Serotonin-norepinephrine reuptake Inhibitor |
| Venlafaxine | Serotonin-norepinephrine reuptake Inhibitor |
| Amitriptyline | Tricyclic antidepressant |
| Imipramine | Tricyclic antidepressant |
| Nortriptyline | Tricyclic antidepressant |
| Trimipramine | Tricyclic antidepressant |
| Clomipramine | Tricyclic antidepressant |
| Dosulepin | Tricyclic antidepressant |
| Maprotiline | Tetracyclic antidepressant |
| Mianserin | Tetracyclic antidepressant |
| Mirtazapine | Tetracyclic antidepressant |
| Tranylcypromine | Monoamine oxidase inhibitor |
| Selegiline | Monoamine oxidase inhibitor |
| Reboxetine | Noradrenaline reuptake inhibitor |

Table S3. List of control agents and drug class

| Control Drugs | Drug Class |
|---|---|
| Pantoprazole | Proton pump inhibitors |
| Clofibrate | Fibrates |
| Rifaximin | Antibiotic |
| Acarbose | Alpha-glucosidase inhibitors |
| Ipriflavone | Isoflavone |

Table S4. Main clinical demographic characteristics of STAR*D

| | |
|---|---|
| Number of individuals N | 1163 |
| Level 1 citalopram remitters | 506 |
| Level 1 citalopram non-remitters | 657 |
| Female ratio | 0.58 |
| Mean age (SD) | 43.33 (13.49) |
| Mean baseline QIDS-C score (SD) | 16.14 (3.16) |

Table S5. Ranking of ADs and control drugs in A375

| Drug | Rank | P-value |
|---|---|---|
| Trimipramine | 2.6 | 0.025 |
| Escitalopram | 2.8 | 0.0296 |
| Maprotiline | 4 | 0.0642 |
| Sertraline | 5.3 | 0.1085 |
| Venlafaxine | 5.4 | 0.1115 |
| Imipramine | 5.5 | 0.1127 |
| Citalopram | 6.6 | 0.1581 |
| Pantoprazole | 10.5 | 0.3373 |
| Clofibrate | 10.8 | 0.3504 |
| Rifaximin | 13.2 | 0.4977 |
| Mirtazapine | 13.5 | 0.5123 |
| Trazodone | 13.5 | 0.5123 |
| Selegiline | 14 | 0.5381 |
| Clomipramine | 14.7 | 0.5777 |
| Nortriptyline | 15.3 | 0.6073 |
| Duloxetine | 15.4 | 0.6135 |
| Reboxetine | 15.4 | 0.6135 |
| Acarbose | 15.9 | 0.6392 |
| Fluoxetine | 16 | 0.6488 |
| Ipriflavone | 17.6 | 0.7162 |
| Amitriptyline | 19.4 | 0.8054 |
| Tranylcypromine | 21.3 | 0.8815 |
| Dosulepin | 21.9 | 0.9000 |
| Paroxetine | 22.7 | 0.9285 |
| Mianserin | 23.5 | 0.9527 |
| Fluvoxamine | 24.2 | 0.9715 |

Note. Citalopram and Escitalopram are highlighted in blue and control drugs are highlighted in yellow

Table S6. Ranking of ADs and control drugs in MCF7

| Drug | Avg Rank | Perm.p.value |
|---|---|---|
| Rifaximin | 1.6 | 0.005 |
| Escitalopram | 2 | 0.0085 |
| Amitriptyline | 3.6 | 0.0477 |
| Venlafaxine | 5.2 | 0.1004 |
| Pantoprazole | 6.4 | 0.1473 |
| Clomipramine | 6.7 | 0.1596 |
| Clofibrate | 7.2 | 0.1885 |
| Fluvoxamine | 8.6 | 0.2492 |
| Sertraline | 9.2 | 0.2804 |
| Maprotiline | 10.1 | 0.3192 |
| Acarbose | 10.2 | 0.3238 |
| Citalopram | 11.1 | 0.3723 |
| Reboxetine | 13.8 | 0.5212 |
| Fluoxetine | 14.4 | 0.5527 |
| Trazodone | 15.2 | 0.6027 |
| Imipramine | 17.2 | 0.6931 |
| Trimipramine | 17.6 | 0.7146 |
| Paroxetine | 18.8 | 0.7742 |
| Ipriflavone | 19.1 | 0.7908 |
| Tranylcypromine | 19.2 | 0.7965 |
| Nortriptyline | 19.2 | 0.7965 |
| Mianserin | 20.2 | 0.8427 |
| Selegiline | 21.6 | 0.9004 |
| Dosulepin | 23.1 | 0.9442 |
| Mirtazapine | 23.8 | 0.9662 |
| Duloxetine | 25.9 | 0.9988 |

Note. Citalopram and Escitalopram are highlighted in blue and control drugs are highlighted in yellow

Table S7. Ranking of ADs and control drugs in PC3

| Drug | Avg Rank | Perm.p.value |
|---|---|---|
| Rifaximin | 1.7 | 0.0062 |
| Ipriflavone | 3.2 | 0.0362 |
| Mirtazapine | 3.5 | 0.0431 |
| Citalopram | 4.1 | 0.0596 |
| Fluvoxamine | 4.2 | 0.0638 |
| Trimipramine | 6 | 0.1327 |
| Paroxetine | 8.3 | 0.2381 |
| Escitalopram | 9.2 | 0.2812 |
| Pantoprazole | 10.6 | 0.3492 |
| Clomipramine | 11.7 | 0.4069 |
| Sertraline | 12.1 | 0.4285 |
| Maprotiline | 12.6 | 0.4581 |
| Reboxetine | 13.6 | 0.5146 |
| Venlafaxine | 13.8 | 0.5288 |
| Nortriptyline | 16 | 0.6331 |
| Amitriptyline | 16.4 | 0.6554 |
| Trazodone | 17.2 | 0.6954 |
| Clofibrate | 17.2 | 0.6954 |
| Acarbose | 17.7 | 0.7196 |
| Imipramine | 18.8 | 0.7723 |
| Dosulepin | 19.2 | 0.7954 |
| Mianserin | 22.2 | 0.9123 |
| Duloxetine | 22.2 | 0.9123 |
| Fluoxetine | 22.5 | 0.9238 |
| Tranylcypromine | 22.8 | 0.9354 |
| Selegiline | 24.2 | 0.9723 |

Note. Citalopram and Escitalopram are highlighted in blue and control drugs are highlighted in yellow

Table S8. Ranking of ADs and control drugs in HA1E

| Drug | Avg Rank | Perm.p.value |
|---|---|---|
| Imipramine | 1.8 | 0.0092 |
| Pantoprazole | 1.8 | 0.0092 |
| Clofibrate | 3.6 | 0.0442 |
| Fluvoxamine | 4.9 | 0.0915 |
| Sertraline | 5.6 | 0.1204 |
| Venlafaxine | 5.6 | 0.1204 |
| Fluoxetine | 8.8 | 0.2569 |
| Rifaximin | 10.5 | 0.3404 |
| Mirtazapine | 11 | 0.3677 |
| Trazodone | 11.2 | 0.3792 |
| Acarbose | 11.7 | 0.4108 |
| Dosulepin | 12 | 0.4219 |
| Tranylcypromine | 12.7 | 0.4623 |
| Trimipramine | 13.1 | 0.4815 |
| Citalopram | 13.8 | 0.5146 |
| Maprotiline | 14.6 | 0.5592 |
| Duloxetine | 14.9 | 0.5738 |
| Escitalopram | 18.5 | 0.7662 |
| Paroxetine | 18.7 | 0.7769 |
| Selegiline | 19.3 | 0.8023 |
| Ipriflavone | 21.2 | 0.8742 |
| Amitriptyline | 22.1 | 0.9088 |
| Mianserin | 22.7 | 0.9323 |
| Nortriptyline | 22.9 | 0.9377 |
| Clomipramine | 23.4 | 0.9523 |
| Reboxetine | 24.6 | 0.9812 |

Note. Citalopram and Escitalopram are highlighted in blue and control drugs are highlighted in yellow

Table S9. Ranking of ADs and control drugs in HT29

| Drug | Avg Rank | Perm.p.value |
|---|---|---|
| Acarbose | 1.4 | 0.0062 |
| Trimipramine | 2.7 | 0.0285 |
| Clofibrate | 3.1 | 0.0377 |
| Ipriflavone | 5.2 | 0.1069 |
| Dosulepin | 5.3 | 0.1108 |
| Rifaximin | 7.1 | 0.1862 |
| Duloxetine | 7.7 | 0.2154 |
| Citalopram | 7.9 | 0.2227 |
| Mirtazapine | 8.5 | 0.2508 |
| Imipramine | 9 | 0.2758 |
| Sertraline | 11.4 | 0.3938 |
| Nortriptyline | 11.7 | 0.4058 |
| Venlafaxine | 13.1 | 0.4738 |
| Fluvoxamine | 13.6 | 0.5035 |
| Maprotiline | 14.4 | 0.5462 |
| Amitriptyline | 16.9 | 0.6892 |
| Reboxetine | 17.3 | 0.7054 |
| Selegiline | 19.7 | 0.8165 |
| Paroxetine | 20.6 | 0.8500 |
| Trazodone | 20.6 | 0.8500 |
| Fluoxetine | 20.7 | 0.8523 |
| Tranylcypromine | 20.9 | 0.8619 |
| Clomipramine | 21 | 0.8662 |
| Escitalopram | 21.9 | 0.8935 |
| Mianserin | 24.1 | 0.9631 |
| Pantoprazole | 25.2 | 0.9908 |

Note. Citalopram and Escitalopram are highlighted in blue and control drugs are highlighted in yellow
